# Inhibition of the polyamine synthesis enzyme ornithine decarboxylase sensitizes triple-negative breast cancer cells to cytotoxic chemotherapy

**DOI:** 10.1101/2020.01.08.899492

**Authors:** Renee C. Geck, Jackson R. Foley, Tracy R. Murray Stewart, John M. Asara, Robert A. Casero, Alex Toker

## Abstract

Treatment of triple-negative breast cancer (TNBC) is limited by a lack of effective molecular targeted therapies. Recent studies have identified metabolic alterations in cancer cells that can be targeted to improve responses to standard-of-care chemotherapy regimens. We found that exposure of TNBC cells to cytotoxic chemotherapy drugs leads to alterations in arginine and polyamine metabolites due to a reduction in the levels and activity of a rate-limiting polyamine biosynthetic enzyme ornithine decarboxylase (ODC). The reduction in ODC was mediated by its negative regulator, antizyme, targeting ODC to the proteasome for degradation. Treatment with the ODC inhibitor DFMO sensitized TNBC cells to chemotherapy, but this was not observed in receptor-positive breast cancer cells. Moreover, TNBC cell lines showed greater sensitivity to single-agent DFMO, and ODC levels were elevated in TNBC patient samples. Alterations in polyamine metabolism in response to chemotherapy, as well as preferential sensitization of TNBC cells to chemotherapy by DFMO, suggest that ODC may be a targetable metabolic vulnerability in TNBC.

## Introduction

In the past two decades, a renewed interest in tumor metabolism has led to the identification of novel therapeutic vulnerabilities in a variety of cancers (1,2). This is especially promising for tumor types that lack effective targeted therapies, such as triple-negative breast cancer (TNBC). TNBC comprises 15-20% of breast cancer cases, but accounts for a disproportionately high percentage of breast cancer-related deaths (3). This is in part due to a lack of targeted therapies for this molecular subtype of breast cancer, and the standard of care treatment for TNBC remains genotoxic chemotherapy. Previous studies have identified metabolic vulnerabilities in breast cancer cells including nucleotide metabolism, glutathione biosynthesis and glutamine metabolism that improve response to chemotherapy drugs (4–6).

The amino acid arginine has been extensively studied in the context of cancer metabolism and has been suggested to contribute to the development and progression of cancer (7). Arginine is involved in numerous cell growth control processes, including protein synthesis, nitric oxide (NO) production and polyamine biosynthesis, as well as cellular energy production via the TCA cycle (8). Furthermore, arginine plays an important role in the immune system, as it is required for full activation of natural killer and T-cells (9,10). A number of studies aimed at targeting arginine metabolism have focused on arginine auxotroph tumors, since they are sensitive to arginine depletion (11,12). However, less than 10% of breast tumors lack the rate-limiting arginine synthesis enzyme argininosuccinate synthase (ASS1) (13). Therefore, additional branches of arginine metabolism may be altered in breast cancer and represent potential metabolic vulnerabilities.

The polyamines putrescine, spermidine and spermine are cationic molecules that are synthesized from arginine following its conversion to ornithine and are essential for eukaryotic cell growth and differentiation (14). Functions attributed to polyamines include binding to nucleic acids and chromatin, stabilizing cellular membranes, regulating ion channels and scavenging free radicals (15). Polyamines can form hydrogen bonds with anions, such as nucleic acids, proteins and phospholipids, and lead to condensation of DNA and chromatin (16,17). Ornithine decarboxylase (ODC), the first rate-limiting enzyme in polyamine synthesis, is regulated transcriptionally, translationally, and post-translationally (18). Moreover, ODC levels and activity are altered in response to extracellular stimuli such as hormones and growth factors, as well as changes in intracellular free polyamine levels (15,18,19).

Elevated polyamine levels have been detected in tumors, including breast tumors, and increased polyamine synthesis has been shown to promote tumor initiation and growth (14,17). Combined inhibition of polyamine uptake and synthesis results in antiproliferative effects in breast cancer cell lines and mouse models, suggesting that targeting polyamine metabolism may be therapeutically effective in breast cancer patients (20,21). Although drugs that target polyamine metabolism have not yet been approved for therapeutic use in cancer, clinical trials using polyamine uptake and synthesis inhibitors are underway for a number of indications, although to date none in breast cancer (22). Here we show that polyamine synthesis is suppressed in response to DNA damaging chemotherapy through the proteasomal degradation of ODC. Further decreasing the polyamine pool using ODC inhibitors increases the sensitivity of TNBC to standard-of-care genotoxic drugs.

## Results

### Genotoxic chemotherapy alters levels of polyamines and related metabolites in TNBC

To evaluate the alterations in intracellular metabolites in breast cancer cells exposed to genotoxic chemotherapy drugs, we measured the levels of polar metabolites in MDA-MB-468 and SUM-159PT TNBC cells following treatment with cisplatin or doxorubicin for 8, 24, or 48 hours. We first determined the concentration of both drugs required for the induction of DNA damage as measured by increased H2AX phosphorylation, while remaining below the IC_50_ for cell death (Fig. 1, *A* and *B*). Based on this analysis, both cell lines were treated with 0.5 µM doxorubicin and 2.5 µM cisplatin, and targeted metabolomics profiling of 186 metabolites was performed using LC-MS/MS (Figs. 1*C* and S1). Increases in pyrimidine nucleotides were observed in response to DNA damage, consistent with previous reports (4).

**Fig. 1:**
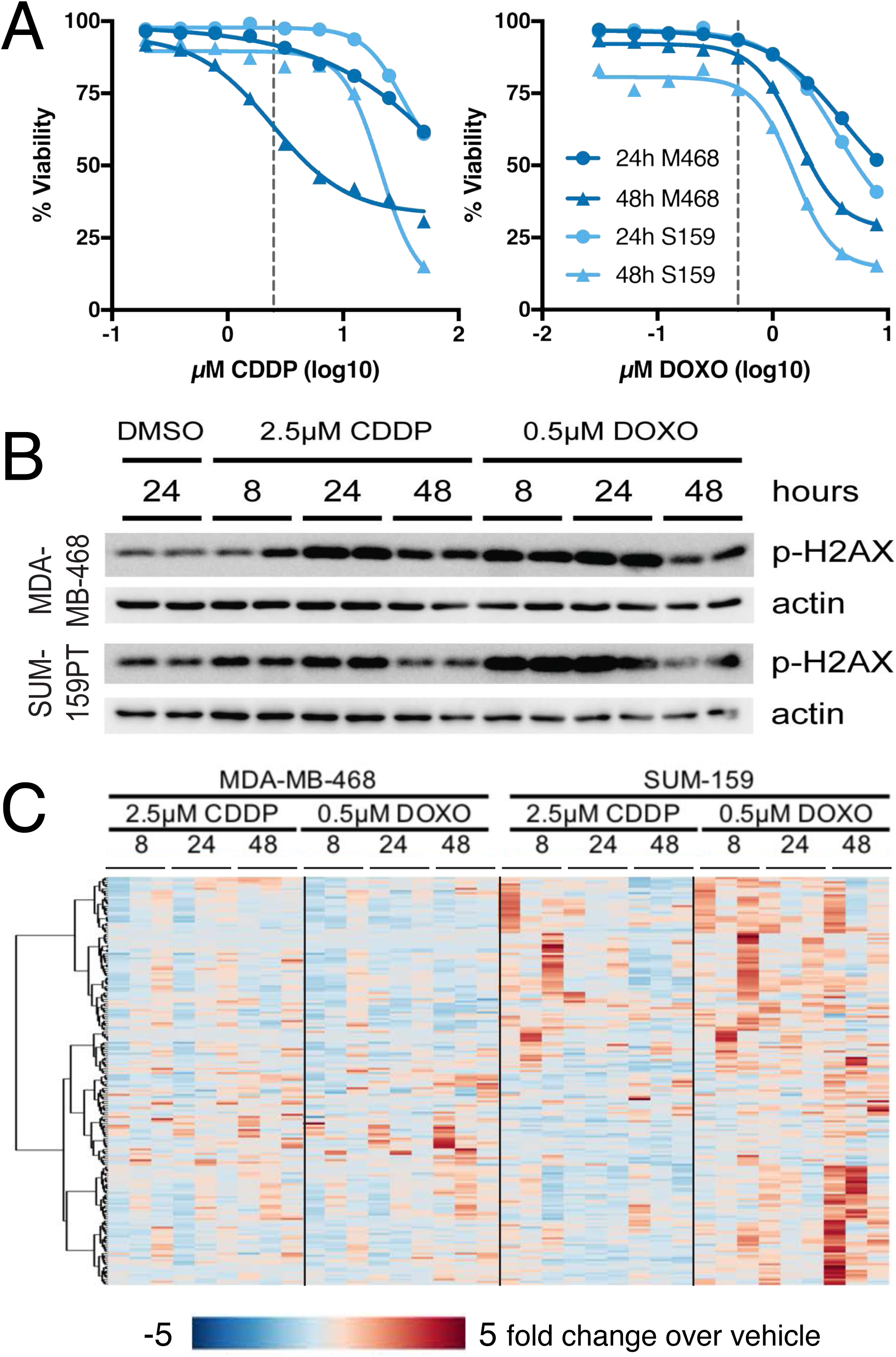
Genotoxic chemotherapy alters TNBC metabolism. (A) Viability measured by propidium iodide uptake following 24 or 48 hours of exposure to chemotherapy agents. Selected concentrations (2.5µM cisplatin, 0.5µM doxorubicin) denoted by dashed lines. Nonlinear curve fit by four parameter logistic regression. (B) Representative immunoblot of phospho-S139 histone H2A.X (p-H2A.X) following exposure to chemotherapy agents at times and doses used for metabolite measurements. (C) Fold change metabolite abundance over 24h vehicle control for 186 metabolites measured by LC-MS/S in MDA-MB-468 and SUM-159PT cells treated with 2.5µM cisplatin (CDDP) or 0.5µM doxorubicin (DOXO).

We focused on metabolites involved in arginine metabolism, including the urea cycle, nitric oxide (NO) cycle, polyamine metabolism, proline metabolism and creatine synthesis (Fig. 2*A*). Twelve metabolites involved in arginine metabolism were detected by our LC-MS/MS platform (Fig. 2, *A* and *B*). In response to genotoxic drugs, the most upregulated metabolite in arginine metabolism was ornithine, and the most decreased was S-adenosyl methionine (SAM) (Fig. 2, *B* and *C*). Ornithine can be synthesized either from arginine by arginase, or from glutamine via ornithine aminotransferase, although previous reports have shown that in transformed cells, ornithine is derived exclusively from arginine (23). In MDA-MB-468 and SUM-159PT TNBC cells, carbons from ^13^C_6_-arginine, but not ^13^C_5_-glutamine, were incorporated into ornithine, even though both exogenous ^13^C_6_-arginine and ^13^C_5_-glutamine efficiently labeled their respective intracellular pools, and this was unaffected in doxorubicin-treated cells (Fig. 2*D*). Since both ornithine and SAM are required for the synthesis of polyamines, but polyamines were not detectable on our LC-MS/MS platform, we measured polyamine levels by conventional HPLC analysis (Fig. 2*E*). After 48 hours of exposure to genotoxic drugs, putrescine and spermidine were significantly decreased by doxorubicin, and also decreased in response to cisplatin. These data suggest that genotoxic drugs alter arginine metabolites involved in polyamine synthesis in TNBC cells.

**Fig. 2:**
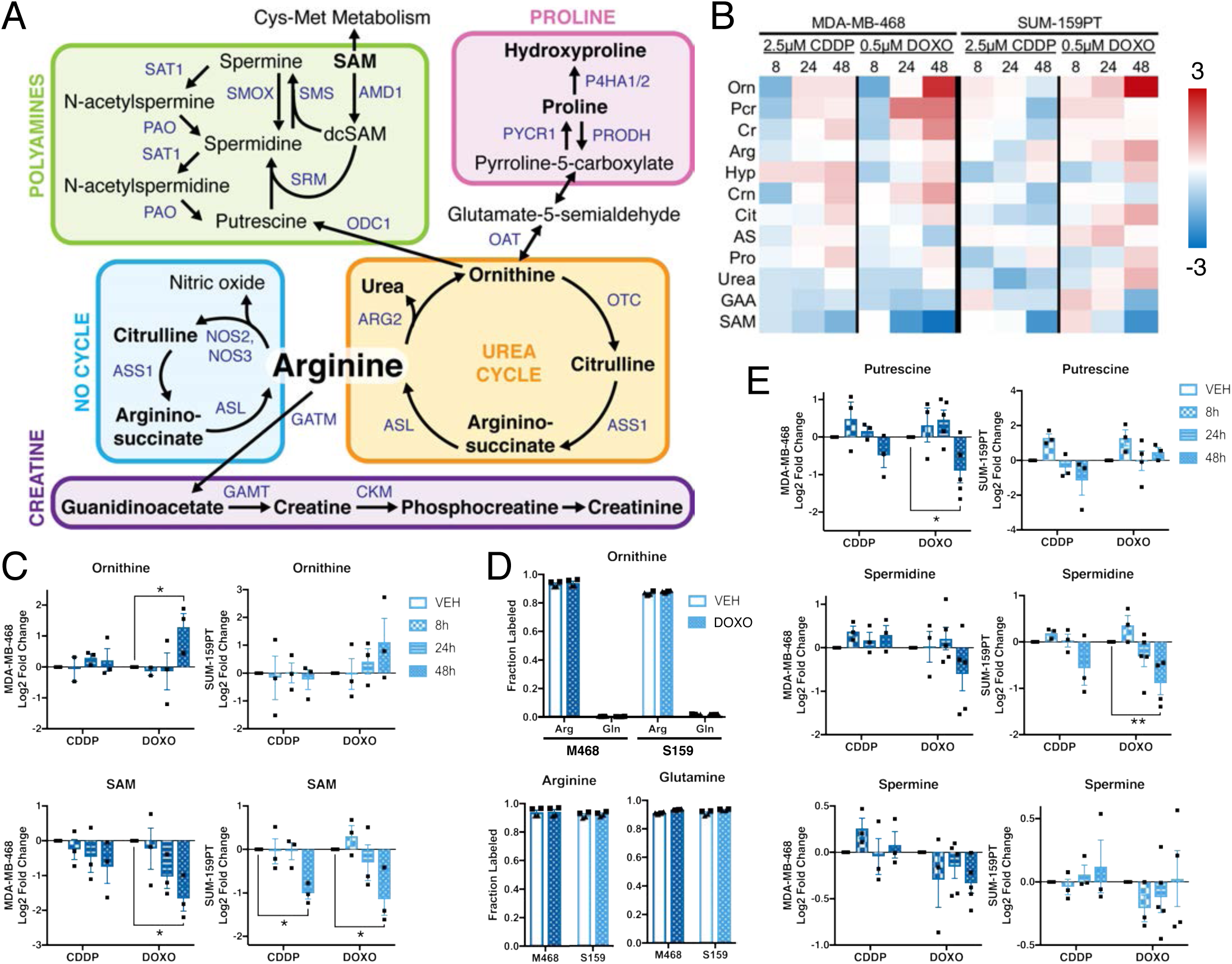
Genotoxic chemotherapy alters arginine and polyamine metabolism. (A) Arginine synthesis and degradation. Metabolites in black, genes in blue. Bold text indicates metabolites detected by LC-MS/MS. (B) Fold change metabolite abundance over 24h vehicle control for arginine metabolites in MDA-MB-468 and SUM-159PT cells treated with 2.5µM cisplatin (CDDP) or 0.5µM doxorubicin (DOXO) (n=3). (C) Relative abundance of ornithine and S-adenosyl methionine (SAM) by LC-MS/MS; vehicle at 24h (n=3). (D) Fraction metabolite pool labeled as measured by LC-MS/MS following incubation with ^13^C_6_-arginine or ^13^C_5_-glutamine in the presence of vehicle or 0.5µM doxorubicin for 48 hours (n=2 biological replicates; technical replicates denoted by shared symbol). (E) Relative abundance of polyamines by HPLC; vehicle control at 24h (n=3-5). All error bars represent SEM. *, P<0.05; **, P<0.01 by 2-way ANOVA. Gene abbreviations: ALDH18A1, Aldehyde Dehydrogenase 18 Family Member A1; AMD1, Adenosylmethionine Decarboxylase 1; ARG2, Arginase 2; ASL, Argininosuccinate Lyase; ASS1, Argininosuccinate Synthase 1; CPS1, Carbamoyl-Phosphate Synthase 1; dcSAM, decarboxylated S-adenosyl methionine; GAMT, Guanidinoacetate N-Methyltransferase; GATM, Glycine Amidinotransferase; NOS, Nitric Oxide Synthase; OAT, Ornithine Aminotransferase; OAZ1, Ornithine Decarboxylase Antizyme 1; ODC1, Ornithine Decarboxylase; OTC, Ornithine Carbamoyltransferase; PAO, Polyamine Oxidase; PRODH, Proline Dehydrogenase 1; PYCR1, Pyrroline-5-Carboxylate Reductase 1; SAM, S-adenosyl methionine; SAT1, Spermidine/Spermine N1-Acetyltransferase 1; SMOX, Spermine Oxidase; SMS, Spermine Synthase; SRM, Spermidine Synthase. Metabolite abbreviations: Arg, L-arginine; AS, L-argininosuccinate; Cit, citrulline; Cr, creatine; Crn, creatinine; GAA, guanidoacetate; Hyp, hydroxyproline; Orn; ornithine; Pcr, phosphocreatine; Pro, proline; Put, putrescine; SAM, S-adenosyl-L-methionine; Spm, spermine; Spd, spermidine.

### Chemotherapy decreases levels and activity of ornithine decarboxylase

We reasoned that the altered levels of polyamine metabolites were due to changes in enzyme levels or activity in response to chemotherapy exposure. ODC catalyzes the first rate-limiting step in polyamine synthesis, specifically the conversion of ornithine to putrescine (Fig. 2*A*). Decreased ODC levels or activity could account for the observed decreases in polyamines and increases in ornithine (Fig. 2, *C* and *E*), though ornithine could also be elevated by increased activity of arginase II (ARG2) (Fig. 2*A*). Cisplatin and doxorubicin increased total ODC protein at early timepoints (8 hours), consistent with a stress response (22), but led to decreased ODC and increased ARG2 at later timepoints (Fig. 3*A*). This pattern of ODC expression was also observed in multiple TNBC and non-TNBC breast cancer cell lines (Fig. 3, *B-D*). By contrast, alterations in ARG2 expression were not significant across all lines. We also confirmed a concomitant decrease in ODC activity after 24 and 48 hours of exposure to cisplatin or doxorubicin (Fig. 4*A*). A significant decrease in putrescine at 72 and 96 hours was also observed, in addition to significantly reduced spermidine after 96 hours of doxorubicin treatment and a time-dependent decreasing trend in spermine concentration (Fig. 4*B*).

**Fig. 3:**
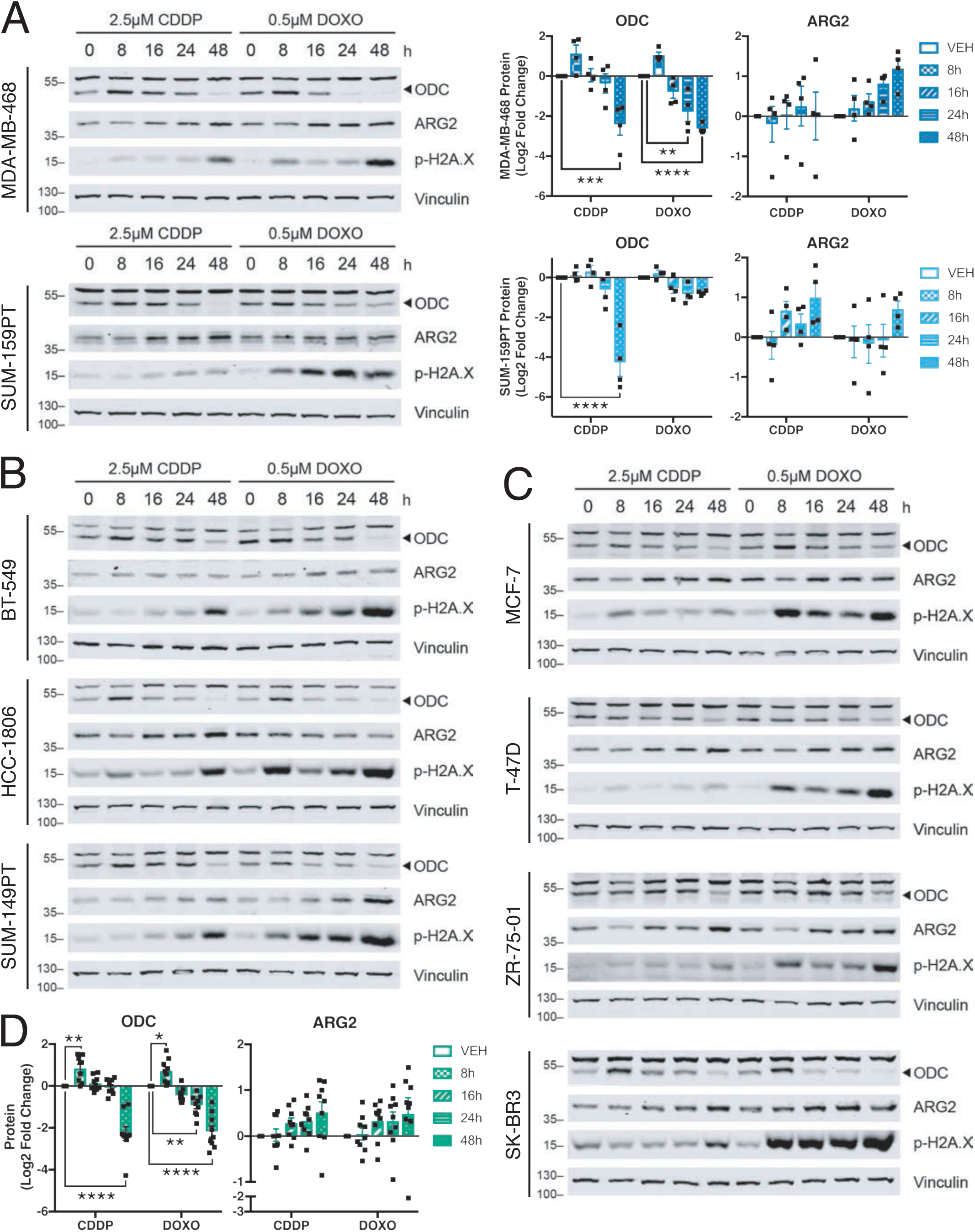
Chemotherapy decreases ODC and increases ARG2 proteins. (A) Representative immunoblots and quantification (n=4) of total ornithine decarboxylase (ODC) and arginase II (ARG2) proteins, and phospho-S139 histone H2A.X (p-H2A.X) following exposure to chemotherapy agents in MDA-MB-468 and SUM-159PT cells, (B) other TNBC cells, and (C) non-TNBC cells. ‘0h’ chemotherapy treatment indicates 24h vehicle control. (D) Average quantification of n=4 immunoblots of ODC and ARG2 from nine breast cancer cell lines in A-C following chemotherapy exposure. Band above 55kD in ODC blots is non-specific. All error bars represent SEM. *, P<0.05; **, P<0.01; ***, P<0.001; ****, P<0.0001 by 2-way ANOVA.

**Fig. 4:**
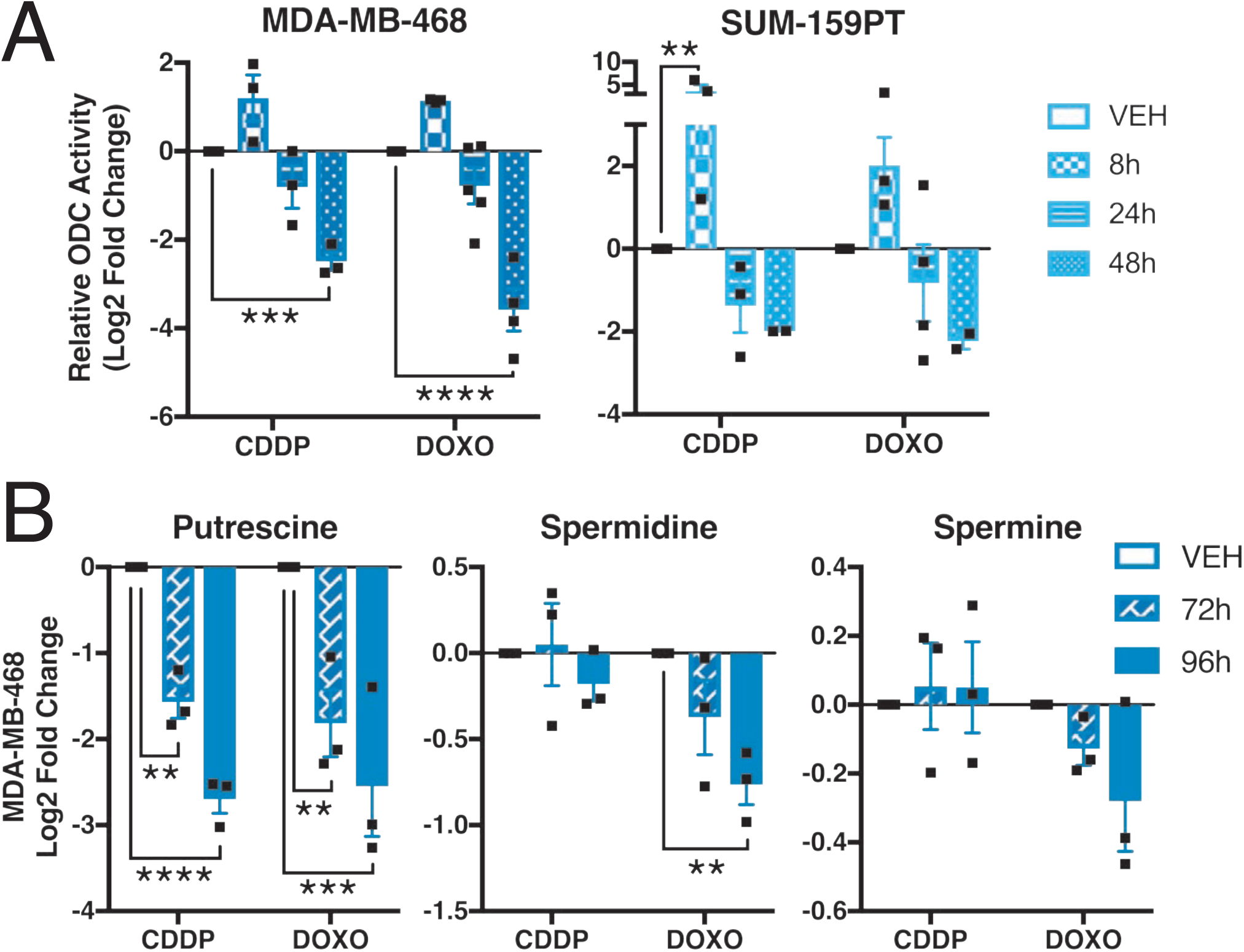
Chemotherapy decreases ODC activity and polyamine levels. (A) ODC activity measured by CO_2_ release following exposure to 2.5µM cisplatin or 0.5µM doxorubicin; vehicle control at 24h (n=3). (B) Relative abundance of polyamines in MDA-MB-468 cells by HPLC; vehicle control at 72h (n=3). All error bars represent SEM. **, P<0.01; ***, P<0.001; ****, P<0.0001 by 2-way ANOVA.

To investigate the mechanism by which ODC protein and activity are decreased following chemotherapy exposure, we first evaluated transcriptional regulation of *ODC1*, which is a well-characterized target of c-Myc (24). Depletion of c-Myc using siRNA did not alter the overall ODC response to chemotherapy (Fig. 5*A*), although *ODC1* transcript was increased following chemotherapy exposure (Fig. 5*B*). ODC protein is post-translationally regulated by the activity of antizyme, which binds ODC to promote its ubiquitin-independent degradation by the 26S proteasome (25). Pre-treatment with the proteasome inhibitor MG132 rescued the decrease in ODC protein following doxorubicin exposure (Fig. 5*C*). siRNA against antizyme decreased the corresponding transcript of *OAZ1* by over 80% (Fig. 5*D*) and blocked the decrease in ODC in response to doxorubicin (Fig. 5*E*). Therefore, it appears that genotoxic drugs decrease polyamines by reducing ODC protein and activity, possibly through the negative regulator antizyme.

**Fig. 5:**
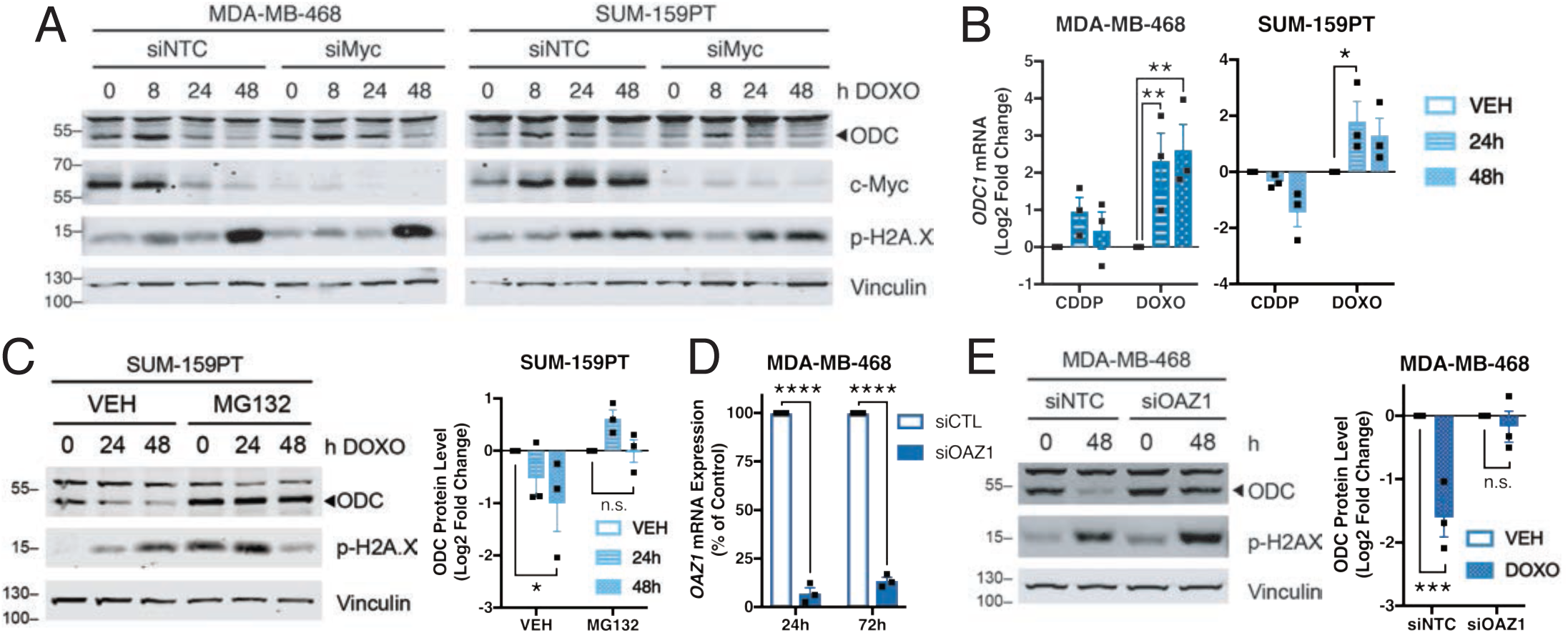
Chemotherapy regulates ODC via proteasomal degradation. (A) Representative immunoblots of ODC and c-Myc in cells pre-treated for 24h with 20nM non-targeting siRNA or siRNA *MYC*, then treated with doxorubicin for indicated times (n=2). (B) qRT-PCR of *ODC1* transcript following treatment with 2.5µM cisplatin or 0.5µM doxorubicin, relative to 24h vehicle control. (C) Representative immunoblots of ODC in SUM-159PT cells pre-treated for 2h with proteasome inhibitor MG132 followed by addition of vehicle or doxorubicin. (n=3) (D) qRT-PCR of *OAZ1* transcript following treatment with 20nM indicated siRNA (n=3). (E) Representative immunoblots and quantification of ODC in MDA-MB-468 cells pre-treated for 24 hours with 20nM siOAZ1 followed by 48 hours addition of vehicle or 0.5µM doxorubicin (n=3). Band above 55kD in ODC blots is non-specific. All error bars represent SEM. *, P<0.05; **, P<0.01; ***, P<0.001 by 2-way ANOVA.

### Targeting polyamine synthesis increases sensitivity to chemotherapy

Polyamines promote cell cycle progression (26), and depletion of ODC or polyamines induces cell cycle arrest at the G2/M phase (27–29), where cells are more sensitive to DNA damage induced by cisplatin and doxorubicin (30–32). Since we observed a decrease in polyamines and ODC activity following chemotherapy treatment, we reasoned that targeting ODC to further decrease polyamines could increase tumor cell killing. Treatment with the irreversible suicide inhibitor of ODC, α-difluoromethylornithine (DFMO), sensitized both MDA-MB-468 and SUM-159PT cells to doxorubicin (Fig. 6*A*). Addition of exogenous putrescine or spermidine did not rescue this sensitization (Fig. 6*B*). A previous study reported that DFMO can decrease colon cancer cell growth by increasing polyamine recycling, leading to a futile cycle that depletes SAM and nucleotides, and that this effect can be rescued by exogenous thymidine (33). However, in TNBC cells, thymidine addition did not rescue the sensitization to doxorubicin by DFMO (Fig. 6*C*). Treatment of MDA-MB-468 cells with the arginase inhibitor Nω-hydroxy-nor-arginine (NOHA) also increased sensitivity to doxorubicin (Fig. 6*D*), consistent with its ability to decrease polyamine levels (34,35).

**Fig. 6:**
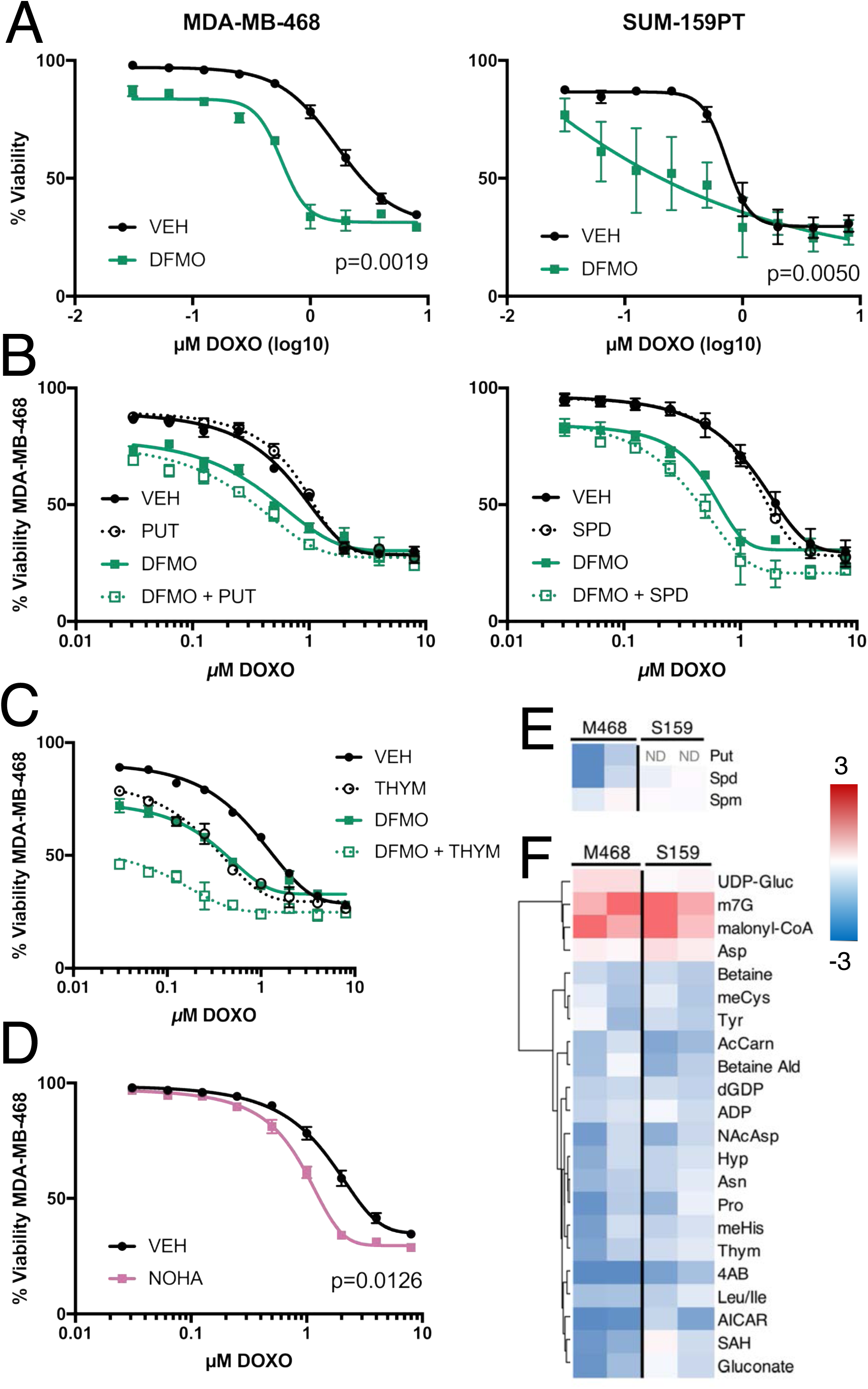
ODC inhibition increases sensitivity to doxorubicin. (A) Viability measured by propidium iodide uptake following 72h pre-treatment with 1mM DFMO or vehicle and 72h exposure to doxorubicin in the presence of DFMO or vehicle (n=3). (B) MDA=MB-468 viability following 72h pretreatment with 1mM DFMO with or without 10µM putrescine or 10µM spermidine and 1mM aminoguanidine, and addition of doxorubicin for 72h (n=2 and n=3). (C) Viability following 72h pretreatment with 1mM DFMO with or without 0.3mM thymidine and addition of doxorubicin for 72h (n=2). (D) Viability following 72h pre-treatment with 0.5mM NOHA and addition of doxorubicin for 72h (n=4) (E) HPLC measurements of polyamines following 72h treatment with 1mM DFMO (n=2). ‘ND’ indicates not detected. (F) Top 10% most significantly altered polar metabolites measured by LC-MS/MS following treatment with 1mM DFMO (n=2). All error bars represent SEM. Nonlinear curve fit by four parameter logistic regression. P-values by unpaired two-tailed t-test. Metabolite abbreviations: 4AB, 4-aminobutyrate; AcCarn, acetylcarnitine-DL; ADP, adenosine diphosphate; AICAR, aminoimidazole carboxamide ribonucleotide; Ald, aldehyde; Asn, asparagine; Asp, aspartate; dGDP, deoxyguanosine diphosphate; Hyp, hydroxyproline; Ile, isoleucine; Leu, leucine; m7G, 7-methylguanosine; meCys, methylcysteine; meHis, 1-methylhistidine; NAcAsp, N-acetyl-L-aspartate; Pro, proline; Put, putrescine; SAH, S-adenosyl-L-homocysteine; Spm, spermine; Spd, spermidine; Thym, thymine; Tyr, tyrosine; UDP-Gluc, uridine diphosphate D-glucose.

To confirm the on-target effects of DFMO, we measured polyamine levels after 72 hours of treatment with DFMO. As expected, putrescine was decreased in MDA-MB-468 cells and undetectable in SUM-159PT cells, and spermidine was also reduced (Fig. 6*E*). To further investigate the effects of DFMO, we measured polar metabolites following exposure to DFMO (Fig. S2). The putrescine metabolite 4-aminobutyrate was significantly decreased, and other metabolites related to nucleotide, one-carbon, and amino acid metabolism were also decreased (Fig. 6*F*). This suggests that these metabolites must be replenished to reverse the effects of DFMO and resulting sensitization to doxorubicin.

Across a panel of breast cancer cell lines representing both luminal A, HER2^+^ and TNBC subtypes, we observed that all TNBC cell lines tested were sensitized to doxorubicin by pretreatment with DFMO, whereas most non-TNBC lines were not (Fig. 7, *A* and *B*). Overall, cell lines that displayed the greatest sensitization to doxorubicin by DFMO are classified as TNBC (Fig. 7*C*). These findings prompted us to determine whether there exists an intrinsic property of TNBC that makes this molecular subtype of breast cancer more dependent on ODC.

**Fig. 7:**
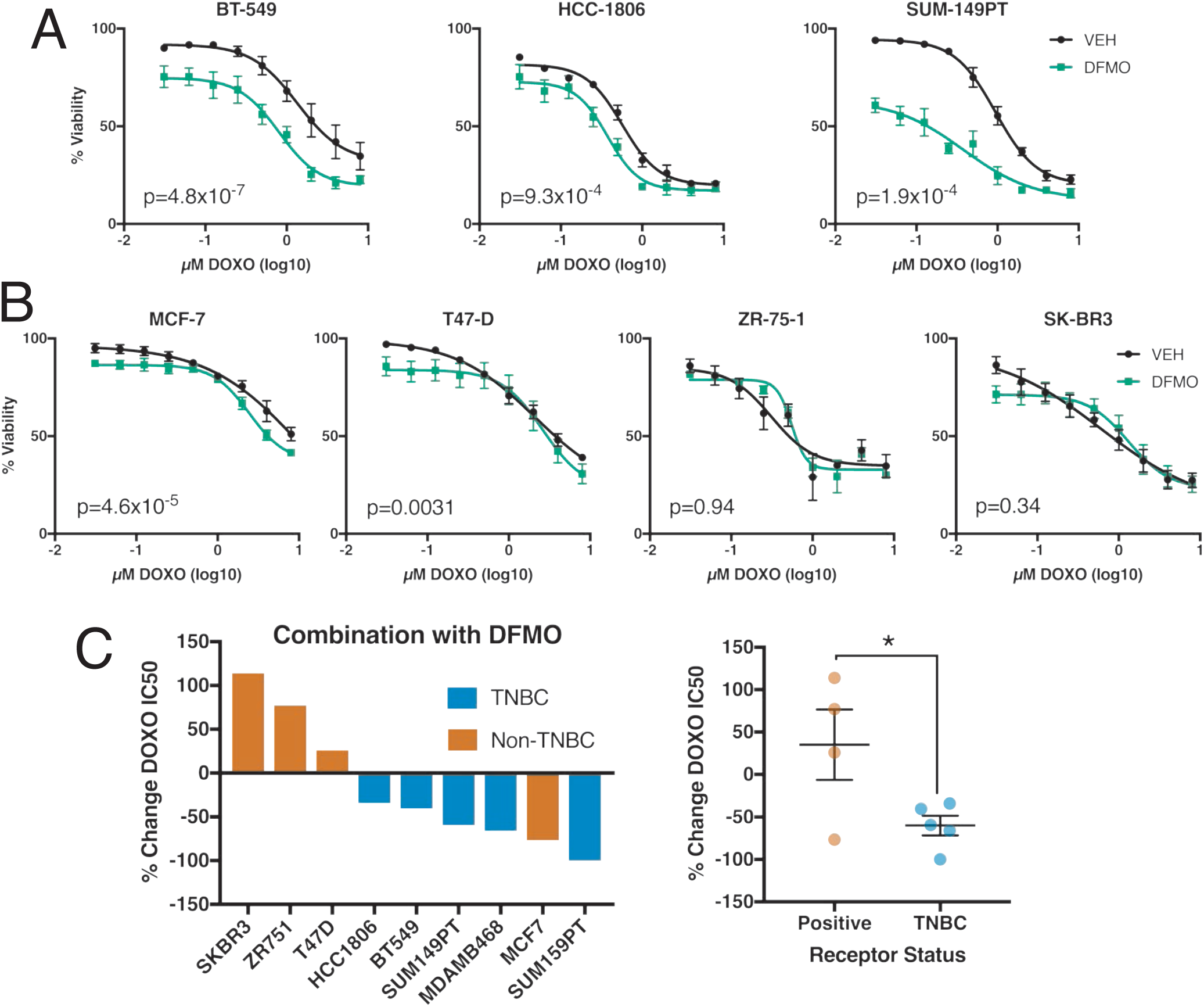
ODC inhibition increases sensitivity of TNBC cells to doxorubicin. (A) Viability measured by propidium iodide uptake following 72h pre-treatment with 1mM DFMO or vehicle and 72h exposure to doxorubicin in the presence of DFMO or vehicle in TNBC and (B) non-TNBC cells (n=3); P-value by paired two-tailed t-test. (C) Percent change doxorubicin IC50 for DFMO over vehicle control from 6A and 7A-B; IC50 calculated by four parameter logistic nonlinear curve fit. All error bars represent SEM. *, P<0.05 by unpaired two-tailed t-test.

### Ornithine decarboxylase is a metabolic vulnerability in TNBC

To investigate the role of arginine and polyamine metabolism in TNBC, we queried the alterations in the transcripts of 1,270 genes in the KEGG ‘metabolic pathways’ (KEGG:hsa01100) in the METABRIC breast cancer dataset (36–38). Of these transcripts, 1,134 were analyzed in 1,904 tumors (Fig. S3). We found that *ODC1* is one of the top 5 most significantly enriched transcripts in TNBC samples (Fig. 8, *A* and *B*). *ODC1* was also enriched in TNBC patient samples from the TCGA provisional breast dataset (Fig. 8*C*). Although we did not observe the same trends for transcript and protein levels of ODC in response to chemotherapy (Fig. 3*A* and Fig. 5*B*), baseline transcript and protein levels positively correlated in the TCGA breast samples for which protein mass spectrometry data is available (Fig. 8*D*). *ODC1* transcript levels were also enriched in the basal subtype of breast cancer, which is commonly associated with TNBC (Fig. 8*E*) (39).

**Fig. 8:**
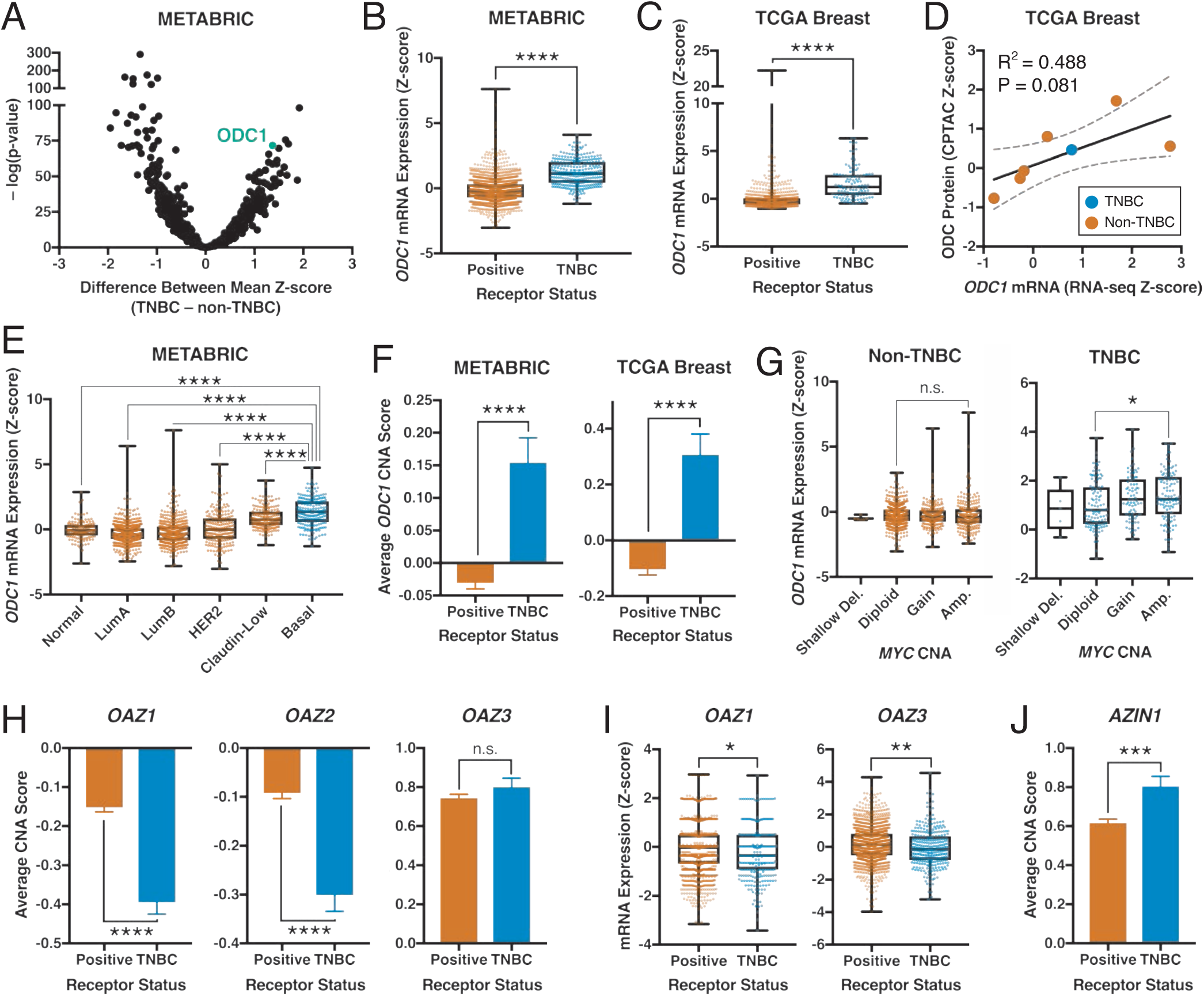
Ornithine decarboxylase levels are decreased in TNBC tumors. (A) Comparison of mRNA z-scores from 1,134 metabolic gene transcripts between 299 TNBC and 1605 non-TNBC patient samples from METABRIC breast cancer dataset. (B) Comparison of *ODC1* mRNA z-scores between METABRIC breast cancer patient samples or (C) TCGA provision breast cancer patient samples, separated by TNBC status. (D) Correlation of *ODC1* mRNA z-scores and ODC protein expression CPTAC z-scores in TCGA provision breast cancer patient samples; dashed lines indicate 90% confidence interval. (E) *ODC1* mRNA z-scores in METABRIC samples grouped by PAM50 subtype. (F) Average copy number alterations of *ODC1* in METABRIC and TCGA provisional breast cancer patient samples scored by type as −1, shallow deletion; 0, diploid; 1, gain; 2, amplification. (G) Comparison of *ODC1* mRNA z-scores between METABRIC breast cancer patient samples separated by TNBC status and grouped by *MYC* copy number alterations. (H) Copy number alterations in METABRIC patient samples for antizyme genes, scored as in F. (I) mRNA z-scores of antizyme transcripts in METABRIC samples grouped by TNBC status. (J) Copy number alterations in METABRIC patient samples for antizyme inhibitor gene *AZIN1*, scored as in F. All error bars represent SEM. *, P<0.05; **, P<0.01 ***, P<0.001; ****, P<0.0001 by Welch’s t-test (A-C,I), one-way ANOVA (E,G), or two-tailed unpaired t-test (F,H,J); linear curve fit, p-value, and R^2^ by linear regression (D).

The increased *ODC1* levels in TNBC could be due to changes in *ODC1* copy number, since the average *ODC1* copy number was higher in TNBC compared to non-TNBC (Fig. 8*F*). *MYC* amplification also correlated with increased *ODC1* transcript in TNBC, but not in non-TNBC patient samples (Fig. 8*G*). We also observed decreased copy number and transcript levels for multiple antizyme genes and transcripts (*OAZ1/2/3*) in TNBC (Fig. 8, *H* and *I*), as well as increased copy number of AZI gene *AZIN1* (Fig. 8*J*) (40).

Enrichment of *ODC1* transcript was also observed in TNBC cells lines according to the Cancer Cell Line Encyclopedia (41) and confirmed by RT-PCR analysis of select breast cancer cell lines (Fig. 9, *A* and *B*), consistent with a previous study of four breast cancer cell lines (42). A trend towards higher baseline protein expression of ODC in TNBC cells was also observed (Fig. 9*C*). Moreover, DFMO treatment significantly reduced proliferation of breast cancer cell lines regardless of their ER/PR/HER2 status (Fig. 9, *D* and *E*). By contrast, DFMO treatment of TNBC cell lines resulted in increased cell death when compared to non-TNBC lines, regardless of doubling time (Fig. 9, *F-H*).

**Fig. 9:**
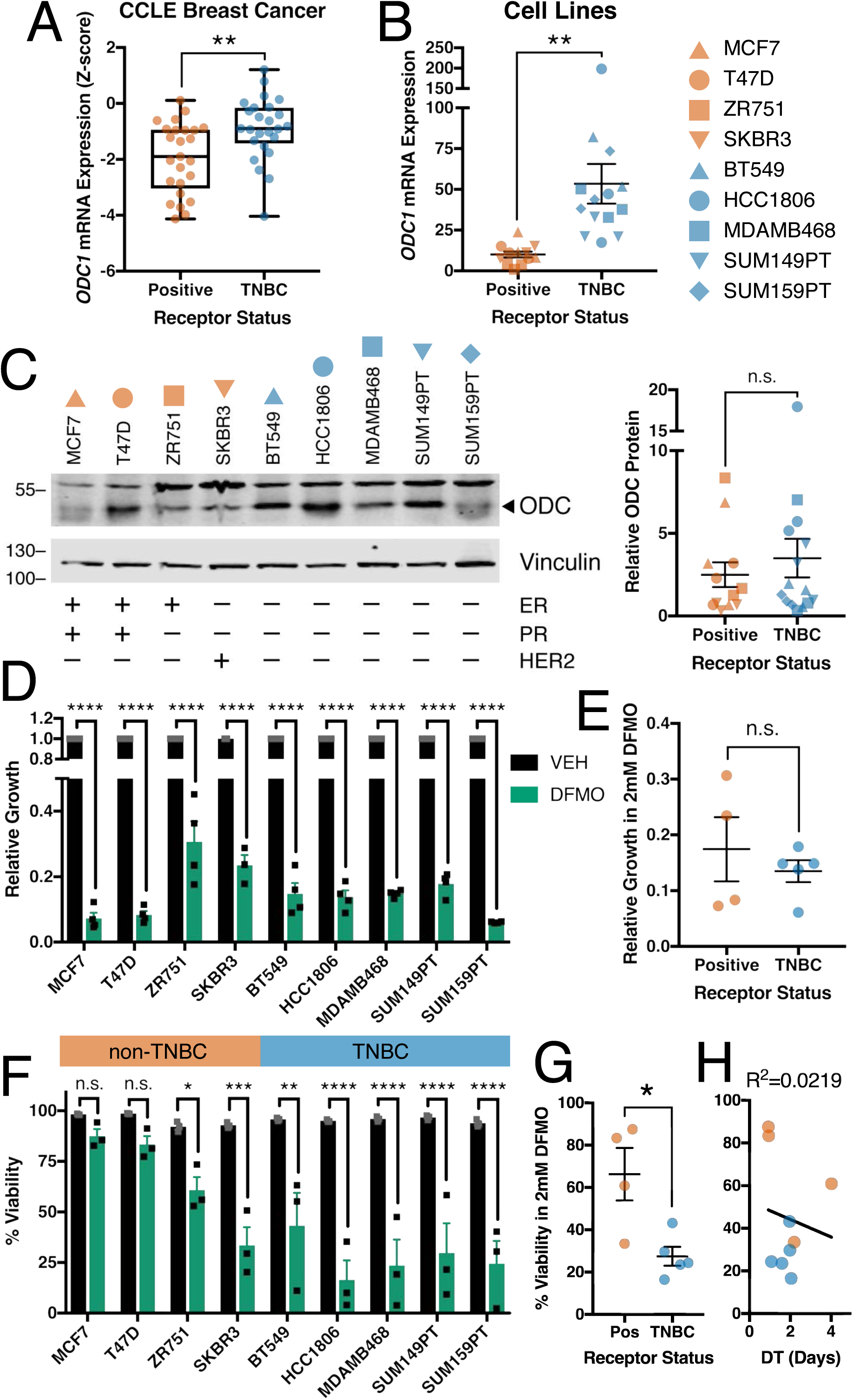
Ornithine decarboxylase is a metabolic vulnerability in TNBC. (A) Comparison of *ODC1* mRNA z-scores in 25 TNBC and 25 non-TNBC breast cancer cell lines Cancer Cell Line Encyclopedia. (B) Comparison of *ODC1* transcript levels measured by qRT-PCR. (C) Representative immunoblots and quantification (n=3) of ODC protein in untreated breast cancer cell lines; band above 55kD in ODC blots is non-specific. (D-E) Growth of breast cancer cell lines measured as population size relative to vehicle after 72h treatment with 2mM DFMO. (F-G) Viability of breast cancer cell lines measured by propidium iodide uptake following 72h treatment with 2mM DFMO. (H) Doubling time of breast cancer cell lines compared to average viability from F; R^2^ by linear regression. All error bars represent SEM. *, P<0.05; **, P<0.01 ***, P<0.001; ****, P<0.0001 by two-tailed unpaired t-test (A-C,E,G) or two-way ANOVA (D,F).

## Discussion

Advances in understanding the reprogramming of metabolic pathways in human cancers have led to the identification of druggable targets that have served as the basis for clinical trials, with the goal of targeting tumor-specific metabolic vulnerabilities. In breast cancer, alterations in glutathione, nucleotide and glutamine metabolism occur in response to genotoxic chemotherapy (4–6). In the present study, we investigated changes in arginine and polyamine-related metabolites in response to genotoxic chemotherapy in TNBC cells. We found that genotoxic drugs decrease ODC levels and activity, with a corresponding reduction in the polyamines putrescine and spermidine. ODC expression is increased in TNBC patient samples and cell lines, to the extent that targeting ODC with the inhibitor DFMO sensitizes TNBC cells to doxorubicin.

TNBC cell lines exposed to chemotherapy showed a significant decrease in the expression and activity of ODC (Fig. 3 and 4). This decrease was not due to alterations in transcript levels nor is it mediated by c-Myc, a major transcriptional regulator of ODC (43). Instead, reduced ODC expression in response to chemotherapy was mediated through degradation by the 26S proteasome (Fig. 5). ODC has a half-life of 10-30 minutes, one of the shortest of any mammalian protein (44). ODC turnover is regulated by binding to antizyme, a protein that binds ODC and targets it for destruction by non-ubiquitin mediated degradation (25). Antizyme itself is translationally regulated through the binding of polyamines to its transcript *OAZ1*, thereby inducing a +1 ribosomal frameshift (25). Thus, antizyme levels are increased when intracellular polyamine concentrations are elevated. The observed initial increases in polyamines after 8 hours of chemotherapy exposure may be sufficient to increase translation of *OAZ1* and subsequently increase antizyme expression (Fig. 3), resulting in ODC degradation. One previous study demonstrated that doxorubicin increases antizyme expression mediated by c-Jun, though the mechanistic basis was not determined (45). An alternative mechanism that could account for ODC regulation is increased antizyme binding to ODC in response to chemotherapy. Moreover, since antizyme inhibitor (AZI) binds antizyme with a greater affinity than ODC, thereby protecting ODC from antizyme-mediated degradation (15,24), decreased AZI would free antizyme to bind ODC. Future studies are necessary to address which of these mechanisms is responsible for regulation of ODC in response to chemotherapy.

Treatment with the ODC inhibitor DFMO sensitizes TNBC cells to chemotherapy drugs, consistent with previous observations that polyamines bind and stabilize DNA and in turn occlude genotoxic agents (46). Conversely, depleting polyamines increases DNA damage and induces cell cycle arrest (47–49). Depletion of ODC or polyamines induces arrest in G2 (28), the cell cycle phase in which tumor cells are most sensitive to damage by cisplatin and doxorubicin (30–32). Depending on the cell type, DFMO arrests cells in either G1 or G2 (27,50–52). Our observations are more consistent with a G2 arrest in response to ODC and DFMO treatment, since this is the phase when cells are most sensitive to genotoxic damage.

Targeting ODC and polyamine synthesis is an attractive antitumor target since high levels of polyamines have been measured in multiple tumor types, and are necessary for transformation and progression (14,17,22). DFMO is approved for treatment of trypanosomiasis and facial hirsutism, and is in clinical trials for treatment or prevention of various tumor types (22). DFMO is of particular chemotherapeutic interest in *MYCN*-amplified neuroblastoma, where Myc-driven overexpression of ODC contributes to tumor hyperproliferation (24). At the time of this study, the only studies of DFMO in combination with genotoxic chemotherapies are in brain tumors, and presently there are no clinical trials in breast cancer using DFMO as single agent therapy or in combination with other drugs. Clinical trials in the late 1990’s established that DFMO reduces polyamines in breast cancer patients, but it did not reduce tumor burden as a single agent (53,54). Application of DFMO in combination with doxorubicin to TNBC models in pre-clinical models would lend further support for the efficacy of this combination, with likely promise due to the apparent low toxicity of DFMO. Combining DFMO with genotoxic chemotherapies could also be effective in other tumor types with elevated ODC activity or expression, including melanoma, esophageal, prostate, and colorectal cancers (28,55–57).

The pronounced effects of DFMO as a single agent on TNBC cell viability (Fig. 9, *F* and *G*) indicate that this molecular subtype of breast cancer is especially reliant on *de novo* polyamine biosynthesis, since in many other cell types, targeting polyamine synthesis is not effective unless combined with a polyamine uptake inhibitor (15). While the specific polyamine transport mechanisms are not well defined (22), it will be interesting to investigate the expression and function of polyamine transporters in TNBC, as deficiencies in transport systems may contribute to a more pronounced reliance on intracellular biosynthesis. Further characterization of polyamine metabolism and the effects of DFMO treatment in TNBC will help identify why this subtype is particularly sensitive.

To our knowledge, this is the first study to profile metabolic changes in response to DFMO in breast cancer. Adding exogenous polyamines or thymidine was insufficient to rescue the effects of DFMO in TNBC (Fig. 6). Importantly, it is evident that treatment of TNBC cells with DFMO also affects metabolites outside of polyamine metabolism, related to nucleotide, one-carbon, and other amino acid metabolism. While in colon tumor cells the effects of DFMO were the result of depleting SAM and nucleotide pools and could be reversed with exogenous thymidine (33), this was not the case in TNBC cells (Fig. 6). Since TNBC cells are more sensitive to DFMO than non-TNBC cells, this implies that an intrinsic genetic or epigenetic property of TNBC renders these tumors more susceptible to ODC inhibition, at least *in vitro*. Whether this represents a metabolic vulnerability in TNBC that can be exploited therapeutically remains to be determined, but given the efficacy of DFMO used in other indications, this warrants exploration in pre-clinical models.

In summary, we have identified changes in arginine and polyamine metabolism in TNBC cells exposed to chemotherapy, due to a decrease of the polyamine synthesis enzyme ODC. By targeting ODC with the inhibitor DFMO, we show that we can sensitize TNBC cells to doxorubicin. Further studies on the mechanisms linking genotoxic damage to ODC degradation, mechanisms of sensitization, and evaluation of combinations in pre-clinical models will provide additional insight for eventual therapeutic applications.

## Experimental procedures

### Cell culture

SUM-159PT and SUM-149PT cells were obtained from Asterand Bioscience; all other cell lines were obtained from the ATCC. Cell lines were authenticated using short tandem repeat profiling, and no cell lines used in this study were found in the database of commonly misidentified cell lines, which is maintained by the International Cell Line Authentication Committee and National Center for Biotechnology Information BioSample. All cells were maintained in RPMI medium (Wisent Bioproducts) containing 10% FBS (Gibco). Cells were passaged for no more than 4 months and routinely assayed for mycoplasma contamination.

### Chemotherapy agents and inhibitors

Doxorubicin was purchased from Cell Signaling Technology and dissolved in DMSO at 10mM; DL-α-Difluoromethylornithine (hydrochloride hydrate) was purchased from Cayman Chemical and dissolved in DMSO at 50mM; spermidine, thymidine, and aminoguanidine hydrochloride were purchased from Sigma and dissolved in water at 10mM,100mM or 1M, respectively; putrescine (dihydrochloride) was purchased from Santa Cruz Biotechnology and dissolved in water at 10mM; N-ω-Hydroxy-L-norarginine acetate salt was purchased from Bachem and dissolved in water at 50mg/mL. Cisplatin was obtained from the Dana Farber Cancer Institute pharmacy at 1mg/mL in PBS. DMSO for use as an organic solvent and vehicle control was purchased from Fisher Scientific.

### Antibodies

ODC (MABS36, 1:100) was purchased from Millipore. ARG2 (ab137069, 1:1000) was purchased from Abcam. pH2A.X^S139^ (9718, 1:1000), beta-actin (4970, 1:1000), c-Myc (5605, 1:1000), and vinculin (13901, 1:1000) were purchased from Cell Signaling Technology. Antibodies were used at indicated dilutions in 5% milk (Andwin Scientific) in TBST buffer (Boston Bioproducts), except for p-H2A.X in 5% BSA (Boston Bioproducts) in TBST.

### LC/MS-MS metabolomics profiling

Cells were maintained in RPMI + 10% FBS, and fresh medium was added at the time cells were treated. For metabolite extraction, medium was aspirated and ice-cold 80% (v/v) methanol was added. Cells and the metabolite-containing supernatants were collected. Insoluble material was pelleted by centrifugation at 20,000 g for 5 minutes. The resulting supernatant was evaporated under nitrogen gas. Samples were resuspended using 20 µL HPLC-grade water for mass spectrometry. For polar metabolite profiling, five microliters from each sample were injected and analyzed using a 5500 QTRAP hybrid triple quadrupole mass spectrometer (AB/SCIEX) coupled to a Prominence UFLC HPLC system (Shimadzu) with HILIC chromatography (Waters Amide XBridge) via selected reaction monitoring (SRM) with polarity switching. A total of 293 endogenous water-soluble metabolites were targeted for steady-state analyses. Electrospray source voltage was +4950 V in positive ion mode and −4500 V in negative ion mode. The dwell time was 3 ms per SRM transition (58). Peak areas from the total ion current for each metabolite were integrated using MultiQuant v2.1.1 software (AB/SCIEX). For ^13^C-labeled experiments, five microliters from each sample (20 uL) were injected with similar methodology as above using a 6500 QTRAP (AB/SCIEX) and integrated using MultiQuant v3.0 software (59). SRMs were created for expected ^13^C incorporation in various forms. Metabolite total ion counts were the integrated total ion current from a single SRM transition and normalized by cellular protein content. Nonhierarchical clustering was performed by Metaboanalyst 4.0 using a Euclidean distance measure and Ward’s clustering algorithm (60).

### Polyamine concentration determinations and ODC activity assays

Cells were washed with PBS and harvested in ODC breaking buffer (25mM Tris-HCl (pH 7.5), 0.1mM EDTA, 2.5mM DTT). Intracellular polyamine concentrations of cell lysates were determined by HPLC following acid extraction and dansylation of the supernatant, as originally described by Kabra et al. (61). Standards prepared for HPLC included diaminoheptane (internal standard), PUT, SPD, and SPM, all of which were purchased from Sigma Chemical Co. (St. Louis, MO, USA). Enzyme activity assays were performed for ODC using radiolabeled substrates, as previously described (62). All enzyme activities and intracellular polyamine concentrations are presented relative to total cellular protein, as determined using Bio-Rad protein dye (Hercules, CA) with interpolation on a bovine serum albumin standard curve.

### Isotope labeling

RPMI powder lacking glutamine, arginine, tryptophan, and glucose was obtained from US Biological Life Sciences and supplemented with 25 µM L-tryptophan (Sigma-Aldrich), 11.1mM D-glucose (Gibco), and 10% dialyzed FBS (Gibco). For arginine labeling, 2mM glutamine (Gibco) and 1.1mM L-arginine-^13^C_6_ hydrochloride (Aldrich) were added; for glutamine labeling, 2mM L-glutamine-^13^C_5_ (Cambridge Isotope Laboratories) and 1.1mM L-arginine hydrochloride (Sigma) were added. Labeled medium, with or without chemotherapy agents, was added to cells, and cellular metabolites were extracted as described above after 48 hours.

### Immunoblotting

Cells were washed with ice-cold PBS (Boston Bioproducts) and lysed in radioimmunoprecipitation buffer (1% NP-40, 0.5% sodium deoxycholate, 0.1% SDS, 150 mM NaCl, 50 mM Tris-HCl (pH 7.5), protease inhibitor cocktail, 50 nM calyculin A, 1mM sodium pyrophosphate, and 20 mM sodium fluoride) for 15 min at 4°C. Cell extracts were cleared by centrifugation at 14,000 rpm for 10 min at 4°C, and protein concentration was measured with the Bio-Rad DC protein assay. Lysates were resolved on acrylamide gels by SDS-PAGE and transferred electrophoretically to nitrocellulose membrane (Bio-Rad) at 100 V for 90 min. Blots were blocked in TBS buffer (10 mmol/L Tris-HCl, pH 8, 150 mmol/L NaCl, Boston Bioproducts) containing 5% (w/v) nonfat dry milk (Andwin Scientific). Membranes were incubated with near-infrared dye-conjugated IRDye 800CW secondary antibodies (LiCor, 1:20,000 in 5% milk-TBST) and imaged on a LiCor Odyssey CLx.

### Quantitative real-time polymerase chain reaction

Total RNA was isolated with the NucleoSpin RNA Plus (MACHEREY-NAGEL) according to the manufacturer’s protocol. Reverse transcription was performed using the TaqMan Reverse Transcription Reagents (Applied Biosciences). Quantitative real-time polymerase chain reaction (qRT-PCR) was performed using a CFX384 Touch Real-Time PCR Detection System (Bio-Rad). Quantification of mRNA expression was calculated by the DCT method with 18S ribosomal RNA as the reference gene.

### RNA interference

siRNA ON-TARGETplus SMARTpools against human ODC1 (L-006668-00), OAZ1 (L-019216-00), and Myc (L-003282-02) were purchased from Dharmacon and dissolved in siRNA buffer (Dharmacon). Cells were transfected with Lipofectamine RNAiMAX Reagent (Invitrogen) in Opti-MEM (Gibco) according to the manufacturer’s protocol (Invitrogen MAN0007825), with a final concentration of 20nM siRNA and 3µL/mL Lipofectamine.

### Propidium iodide viability assay

Cell viability was assayed with a propidium iodide-based plate-reader assay, as previously described (63). Briefly, cells in 96-well plates were treated with a final concentration of 30mM propidium iodide (Cayman Chemical) for 20 min at 37°C. The initial fluorescence intensity was measured in a GENios FL (Tecan) at 560nm excitation/635nm emission. Digitonin (Millipore) was then added to each well at a final concentration of 600mM. After incubating for 20 min at 37°C, the final fluorescence intensity was measured. The fraction of dead cells was calculated by dividing the background-corrected initial fluorescence intensity by the final fluorescence intensity. Viability was calculated by (1 – fraction of dead cells).

### Sulforhodamine B growth assay

Population size as cell confluency was assayed by sulforhodamine B staining, as previously described (64). Briefly, cells in 96-well plates were fixed in 8.3% final concentration of tricarboxylic acid of 1 hour at 4°C, washed three times with water, and stained for 30 minutes with 0.5% sulforhodamine B in 1% acetic acid. Plates were washed 3 times with 1% acetic acid and dye was solubilized in 10mM Tris pH 10.5 before measuring absorbance at 510nm on an Epoch plate reader (BioTek). Relative growth was determined as (treatment absorbance / control absorbance) following 72 hours of growth. Doubling time was determined by fitting an exponential growth equation to population sizes over four days.

### Analysis of public data

Data from METABRIC and CCLE were accessed through cBioPortal, www.cbioportal.org. Data was analyzed using MATLAB R2017b and Prism 7.

### Statistics and reproducibility

Sample sizes, reproducibility, and statistical tests for each figure are denoted in the figure legends. All replicates are biological unless otherwise noted. All error bars represent SEM, and significance between conditions is denoted as *, P<0.05; **, P<0.01; ***, P<0.001; and ****, P<0.0001.

## Acknowledgments

We thank B. Manning (Harvard School of Public Health) for suggestions and critical reagents; J. Brugge (Harvard Medical School), N. Kalaany (Boston Children’s Hospital), and members of the Toker lab (Beth Israel Deaconess Medical Center) for discussion; M. Yuan and C. Dibble (Beth Israel Deaconess Medical Center) for assistance with mass spectrometry.

## Author contributions

R.C.G. and A.T. designed the study and interpreted the results. R.C.G performed all experiments unless otherwise noted. J.R.F. conducted HPLC measurement of polyamines and T.R.M.S. conducted ODC activity assays, both under the direction of R.A.C. J.M.A. assisted with the LC-MS/MS metabolomic studies and data analysis. R.C.G. and A.T. wrote the manuscript.

## Abbreviations used are

ARG2: Arginase 2
ASS1: Argininosuccinate Synthase
AZI: antizyme inhibitor (protein)
AZIN: antizyme inhibitor (gene/transcript)
CDDP: cisplatin
DFMO: α-difluoromethylornithine
DOXO: doxorubicin
NOHA: Nω-hydroxy-nor-arginine
OAZ1: antizyme 1 (gene/transcript)
ODC: Ornithine Decarboxylase
PBT: polyamine blocking therapy
SAM: S-adenosyl methionine
Arg: L-arginine
Orn: ornithine
Put: putrescine
SAM: S-adenosyl-L-methionine
Spm: spermine
Spd: spermidine
TNBC: triple-negative breast cancer

## References

1. Vander Heiden, M. G., and DeBerardinis, R. J. (2017) Understanding the Intersections between Metabolism and Cancer Biology. Cell 168, 657–669

2. DeBerardinis, R. J., and Chandel, N. S. (2016) Fundamentals of cancer metabolism. Science Advances 2, e1600200–e1600200

3. Boyle, P. (2012) Triple-negative breast cancer: epidemiological considerations and recommendations. Ann Oncol 23 Suppl 6, vi7–12

4. Brown, K. K., Spinelli, J. B., Asara, J. M., and Toker, A. (2017) Adaptive reprogramming of de novo pyrimidine synthesis is a metabolic vulnerability in triple-negative breast cancer. Cancer Discovery 7, 391–399

5. Lien, E. C., Lyssiotis, C. A., Juvekar, A., Hu, H., Asara, J. M., Cantley, L. C., and Toker, A. (2016) Glutathione biosynthesis is a metabolic vulnerability in PI(3)K/Akt-driven breast cancer. Nature Cell Biology 18, 572–578

6. Gross, M. I., Demo, S. D., Dennison, J. B., Chen, L., Chernov-Rogan, T., Goyal, B., Janes, J. R., Laidig, G. J., Lewis, E. R., Li, J., Mackinnon, A. L., Parlati, F., Rodriguez, M. L. M., Shwonek, P. J., Sjogren, E. B., Stanton, T. F., Wang, T., Yang, J., Zhao, F., and Bennett, M. K. (2014) Antitumor activity of the glutaminase inhibitor CB-839 in triple-negative breast cancer. Molecular Cancer Therapeutics 13, 890–901

7. Szefel, J., Danielak, A., and Kruszewski, W. J. (2018) Metabolic pathways of L-arginine and therapeutic consequences in tumors. Adv Med Sci 64, 104–110

8. Morris, S. M. (2009) Recent advances in arginine metabolism: roles and regulation of the arginases. British Journal of Pharmacology 157, 922–930

9. Werner, A., Amann, E., Schnitzius, V., Habermeier, A., Luckner-Minden, C., Leuchtner, N., Rupp, J., Closs, E. I., and Munder, M. (2016) Induced arginine transport via cationic amino acid transporter-1 is necessary for human T-cell proliferation. European Journal of Immunology 46, 92–103

10. Hoechst, B., Voigtlaender, T., Ormandy, L., Gamrekelashvili, J., Zhao, F., Wedemeyer, H., Lehner, F., Manns, M. P., Greten, T. F., and Korangy, F. (2009) Myeloid derived suppressor cells inhibit natural killer cells in patients with hepatocellular carcinoma via the NKp30 receptor. Hepatology 50, 799–807

11. Jahani, M., Noroznezhad, F., and Mansouri, K. (2018) Arginine: Challenges and opportunities of this two-faced molecule in cancer therapy. Biomed Pharmacother 102, 594–601

12. Qiu, F., Huang, J., and Sui, M. (2015) Targeting arginine metabolism pathway to treat arginine-dependent cancers. Cancer Letters 364, 1–7

13. Dillon, B. J., Prieto, V. G., Curley, S. A., Ensor, C. M., Holtsberg, F. W., Bomalaski, J. S., and Clark, M. A. (2004) Incidence and distribution of argininosuccinate synthetase deficiency in human cancers: a method for identifying cancers sensitive to arginine deprivation. Cancer 100, 826–833

14. Agostinelli, E., Marques, M. P. M., Calheiros, R., Gil, F. P. S. C., Tempera, G., Viceconte, N., Battaglia, V., Grancara, S., and Toninello, A. (2010) Polyamines: Fundamental characters in chemistry and biology. Amino Acids 38, 393–403

15. Murray-Stewart, T. R., Woster, P. M., and Casero, R. A., Jr. (2016) Targeting polyamine metabolism for cancer therapy and prevention. Biochem J 473, 2937–2953

16. Battaglia, V., DeStefano Shields, C., Murray-Stewart, T., and Casero, R. A., Jr. (2014) Polyamine catabolism in carcinogenesis: potential targets for chemotherapy and chemoprevention. Amino Acids 46, 511–519

17. Nowotarski, S. L., Woster, P. M., and Casero, R. A., Jr. (2013) Polyamines and cancer: implications for chemotherapy and chemoprevention. Expert Rev Mol Med 15, e3

18. Perez-Leal, O., and Merali, S. (2012) Regulation of polyamine metabolism by translational control. Amino Acids 42, 611–617

19. Murray Stewart, T., Dunston, T. T., Woster, P. M., and Casero, R. A., Jr. (2018) Polyamine catabolism and oxidative damage. J Biol Chem 293, 18736–18745

20. Burns, M. R., Graminski, G. F., Weeks, R. S., Chen, Y., and O’Brien, T. G. (2009) Lipophilic lysine-spermine conjugates are potent polyamine transport inhibitors for use in combination with a polyamine biosynthesis inhibitor. J Med Chem 52, 1983–1993

21. Hayes, C. S., Shicora, A. C., Keough, M. P., Snook, A. E., Burns, M. R., and Gilmour, S. K. (2014) Polyamine-blocking therapy reverses immunosuppression in the tumor microenvironment. Cancer Immunol Res 2, 274–285

22. Casero, R. A., Jr., Murray Stewart, T., and Pegg, A. E. (2018) Polyamine metabolism and cancer: treatments, challenges and opportunities. Nat Rev Cancer 18, 681–695

23. Roci, I., Watrous, J. D., Lagerborg, K. A., Lafranchi, L., Lindqvist, A., Jain, M., and Nilsson, R. (2019) Mapping metabolic events in the cancer cell cycle reveals arginine catabolism in the committed SG2M phase. Cell Reports 26, 1691–1700

24. Bachmann, A. S., and Geerts, D. (2018) Polyamine synthesis as a target of MYC oncogenes. J Biol Chem 293, 18757–18769

25. Kahana, C. (2018) The antizyme family for regulating polyamines. J Biol Chem 293, 18730–18735

26. Oredsson, S. M. (2003) Polyamine dependence of normal cell-cycle progression. Biochem Soc Trans 31, 366–370

27. Weicht, R. R., Schultz, C. R., Geerts, D., Uhl, K. L., and Bachmann, A. S. (2018) Polyamine Biosynthetic Pathway as a Drug Target for Osteosarcoma Therapy. Med Sci (Basel) 6

28. He, W., Roh, E., Yao, K., Liu, K., Meng, X., Liu, F., Wang, P., Bode, A. M., and Dong, Z. (2017) Targeting ornithine decarboxylase (ODC) inhibits esophageal squamous cell carcinoma progression. NPJ Precis Oncol 1, 13

29. Anehus, S., Pohjanpelto, P., Baldetorp, B., Langstrom, E., and Heby, O. (1984) Polyamine starvation prolongs the S and G2 phases of polyamine-dependent (arginase-deficient) CHO cells. Mol Cell Biol 4, 915–922

30. Mueller, S., Schittenhelm, M., Honecker, F., Malenke, E., Lauber, K., Wesselborg, S., Hartmann, J. T., Bokemeyer, C., and Mayer, F. (2006) Cell-cycle progression and response of germ cell tumors to cisplatin in vitro. Int J Oncol 29, 471–479

31. Potter, A. J., Gollahon, K. A., Palanca, B. J., Harbert, M. J., Choi, Y. M., Moskovitz, A. H., Potter, J. D., and Rabinovitch, P. S. (2002) Flow cytometric analysis of the cell cycle phase specificity of DNA damage induced by radiation, hydrogen peroxide and doxorubicin. Carcinogenesis 23, 389–401

32. Ling, Y. H., el-Naggar, A. K., Priebe, W., and Perez-Soler, R. (1996) Cell cycle-dependent cytotoxicity, G2/M phase arrest, and disruption of p34cdc2/cyclin B1 activity induced by doxorubicin in synchronized P388 cells. Mol Pharmacol 49, 832–841

33. Witherspoon, M., Chen, Q., Kopelovich, L., Gross, S. S., and Lipkin, S. M. (2013) Unbiased metabolite profiling indicates that a diminished thymidine pool is the underlying mechanism of colon cancer chemoprevention by alpha-difluoromethylornithine. Cancer Discov 3, 1072–1081

34. Singh, R., Pervin, S., Wu, G., and Chaudhuri, G. (2001) Activation of caspase-3 activity and apoptosis in MDA-MB-468 cells by N(omega)-hydroxy-L-arginine, an inhibitor of arginase, is not solely dependent on reduction in intracellular polyamines. Carcinogenesis 22, 1863–1869

35. Singh, R., Pervin, S., Karimi, A., Cederbaum, S., and Chaudhuri, G. (2000) Arginase activity in human breast cancer cell lines: N(omega)-hydroxy-L-arginine selectively inhibits cell proliferation and induces apoptosis in MDA-MB-468 cells. Cancer Research 60, 3305–3312

36. Cerami, E., Gao, J., Dogrusoz, U., Gross, B. E., Sumer, S. O., Aksoy, B. A., Jacobsen, A., Byrne, C. J., Heuer, M. L., Larsson, E., Antipin, Y., Reva, B., Goldberg, A. P., Sander, C., and Schultz, N. (2012) The cBio cancer genomics portal: an open platform for exploring multidimensional cancer genomics data. Cancer Discov 2, 401–404

37. Gao, J., Aksoy, B. A., Dogrusoz, U., Dresdner, G., Gross, B., Sumer, S. O., Sun, Y., Jacobsen, A., Sinha, R., Larsson, E., Cerami, E., Sander, C., and Schultz, N. (2013) Integrative analysis of complex cancer genomics and clinical profiles using the cBioPortal. Sci Signal 6, pl1

38. Pereira, B., Chin, S. F., Rueda, O. M., Vollan, H. K., Provenzano, E., Bardwell, H. A., Pugh, M., Jones, L., Russell, R., Sammut, S. J., Tsui, D. W., Liu, B., Dawson, S. J., Abraham, J., Northen, H., Peden, J. F., Mukherjee, A., Turashvili, G., Green, A. R., McKinney, S., Oloumi, A., Shah, S., Rosenfeld, N., Murphy, L., Bentley, D. R., Ellis, I. O., Purushotham, A., Pinder, S. E., Borresen-Dale, A. L., Earl, H. M., Pharoah, P. D., Ross, M. T., Aparicio, S., and Caldas, C. (2016) The somatic mutation profiles of 2,433 breast cancers refines their genomic and transcriptomic landscapes. Nat Commun 7, 11479

39. Mayer, I. A., Abramson, V. G., Lehmann, B. D., and Pietenpol, J. A. (2014) New Strategies for Triple-Negative Breast Cancer–Deciphering the Heterogeneity. Clinical Cancer Research 20, 782–790

40. Keren-Paz, A., Bercovich, Z., and Kahana, C. (2007) Antizyme inhibitor: a defective ornithine decarboxylase or a physiological regulator of polyamine biosynthesis and cellular proliferation. Biochem Soc Trans 35, 311–313

41. Barretina, J., Caponigro, G., Stransky, N., Venkatesan, K., Margolin, A. A., Kim, S., Wilson, C. J., Lehar, J., Kryukov, G. V., Sonkin, D., Reddy, A., Liu, M., Murray, L., Berger, M. F., Monahan, J. E., Morais, P., Meltzer, J., Korejwa, A., Jane-Valbuena, J., Mapa, F. A., Thibault, J., Bric-Furlong, E., Raman, P., Shipway, A., Engels, I. H., Cheng, J., Yu, G. K., Yu, J., Aspesi, P., Jr., de Silva, M., Jagtap, K., Jones, M. D., Wang, L., Hatton, C., Palescandolo, E., Gupta, S., Mahan, S., Sougnez, C., Onofrio, R. C., Liefeld, T., MacConaill, L., Winckler, W., Reich, M., Li, N., Mesirov, J. P., Gabriel, S. B., Getz, G., Ardlie, K., Chan, V., Myer, V. E., Weber, B. L., Porter, J., Warmuth, M., Finan, P., Harris, J. L., Meyerson, M., Golub, T. R., Morrissey, M. P., Sellers, W. R., Schlegel, R., and Garraway, L. A. (2012) The Cancer Cell Line Encyclopedia enables predictive modelling of anticancer drug sensitivity. Nature 483, 603–607

42. Thomas, T., Kiang, D. T., Janne, O. A., and Thomas, T. J. (1991) Variations in amplification and expression of the ornithine decarboxylase gene in human breast cancer cells. Breast Cancer Res Treat 19, 257–267

43. Bello-Fernandez, C., Packham, G., and Cleveland, J. L. (1993) The ornithine decarboxylase gene is a transcriptional target of c-Myc. Proc Natl Acad Sci U S A 90, 7804–7808

44. Casero, R. A., Jr., and Marton, L. J. (2007) Targeting polyamine metabolism and function in cancer and other hyperproliferative diseases. Nat Rev Drug Discov 6, 373–390

45. Dulloo, I., Gopalan, G., Melino, G., and Sabapathy, K. (2010) The antiapoptotic DeltaNp73 is degraded in a c-Jun-dependent manner upon genotoxic stress through the antizyme-mediated pathway. Proc Natl Acad Sci U S A 107, 4902–4907

46. Nayvelt, I., Hyvönen, M. T., Alhonen, L., Pandya, I., Thomas, T., Khomutov, A. R., Vepsäläinen, J., Patel, R., Keinänen, T. A., and Thomas, T. J. (2010) DNA condensation by chiral α-methylated polyamine analogues and protection of cellular DNA from oxidative damage. Biomacromolecules 11, 97–105

47. Zahedi, K., Bissler, J. J., Wang, Z., Josyula, A., Lu, L., Diegelman, P., Kisiel, N., Porter, C. W., Soleimani, M., N, K., Cw, P., and N, S. M. S. (2007) Spermidine / spermine N 1 - acetyltransferase overexpression in kidney epithelial cells disrupts polyamine homeostasis, leads to DNA damage, and causes G2 arrest. October 45267, 1204–1215

48. Johansson, V. M., Oredsson, S. M., and Alm, K. (2008) Polyamine depletion with two different polyamine analogues causes DNA damage in human breast cancer cell lines. DNA and Cell Biology 27, 511–516

49. Courdi, A., Milano, G., Bouclier, M., and Lalanne, C. M. (1986) Radiosensitization of human tumor cells by alpha-difluoromethylornithine. Int J Cancer 38, 103–107

50. Seidenfeld, J., Block, A. L., Komar, K. A., and Naujokas, M. F. (1986) Altered cell cycle phase distributions in cultured human carcinoma cells partially depleted of polyamines by treatment with difluoromethylornithine. Cancer Res 46, 47–53

51. Saunders, L. R., and Verdin, E. (2006) Ornithine decarboxylase activity in tumor cell lines correlates with sensitivity to cell death induced by histone deacetylase inhibitors. Mol Cancer Ther 5, 2777–2785

52. Yamashita, T., Nishimura, K., Saiki, R., Okudaira, H., Tome, M., Higashi, K., Nakamura, M., Terui, Y., Fujiwara, K., Kashiwagi, K., and Igarashi, K. (2013) Role of polyamines at the G1/S boundary and G2/M phase of the cell cycle. Int J Biochem Cell Biol 45, 1042–1050

53. O’Shaughnessy, J. A., Demers, L. M., Jones, S. E., Arseneau, J., Khandelwal, P., George, T., Gersh, R., Mauger, D., and Manni, A. (1999) Alpha-difluoromethylornithine as treatment for metastatic breast cancer patients. Clin Cancer Res 5, 3438–3444

54. Fabian, C. J., Kimler, B. F., Brady, D. A., Mayo, M. S., Chang, C. H., Ferraro, J. A., Zalles, C. M., Stanton, A. L., Masood, S., Grizzle, W. E., Boyd, N. F., Arneson, D. W., and Johnson, K. A. (2002) A phase II breast cancer chemoprevention trial of oral alpha-difluoromethylornithine: breast tissue, imaging, and serum and urine biomarkers. Clin Cancer Res 8, 3105–3117

55. Yang, D., Hayashi, H., Takii, T., Mizutani, Y., Inukai, Y., and Onozaki, K. (1997) Interleukin-1-induced growth inhibition of human melanoma cells. Interleukin-1-induced antizyme expression is responsible for ornithine decarboxylase activity down-regulation. J Biol Chem 272, 3376–3383

56. Young, L., Salomon, R., Au, W., Allan, C., Russell, P., and Dong, Q. (2006) Ornithine decarboxylase (ODC) expression pattern in human prostate tissues and ODC transgenic mice. J Histochem Cytochem 54, 223–229

57. Hu, H. Y., Liu, X. X., Jiang, C. Y., Lu, Y., Liu, S. L., Bian, J. F., Wang, X. M., Geng, Z., Zhang, Y., and Zhang, B. (2005) Ornithine decarboxylase gene is overexpressed in colorectal carcinoma. World J Gastroenterol 11, 2244–2248

58. Yuan, M., Breitkopf, S. B., Yang, X., and Asara, J. M. (2012) A positive/negative ion-switching, targeted mass spectrometry-based metabolomics platform for bodily fluids, cells, and fresh and fixed tissue. Nat Protoc 7, 872–881

59. Yuan, M., Kremer, D. M., Huang, H., Breitkopf, S. B., Ben-Sahra, I., Manning, B. D., Lyssiotis, C. A., and Asara, J. M. (2019) Ex vivo and in vivo stable isotope labelling of central carbon metabolism and related pathways with analysis by LC-MS/MS. Nat Protoc 14, 313–330

60. Chong, J., Soufan, O., Li, C., Caraus, I., Li, S., Bourque, G., Wishart, D. S., and Xia, J. (2018) MetaboAnalyst 4.0: towards more transparent and integrative metabolomics analysis. Nucleic Acids Res 46, W486–W494

61. Kabra, P. M., Lee, H. K., Lubich, W. P., and Marton, L. J. (1986) Solid-phase extraction and determination of dansyl derivatives of unconjugated and acetylated polyamines by reversed-phase liquid chromatography: improved separation systems for polyamines in cerebrospinal fluid, urine and tissue. J Chromatogr 380, 19–32

62. Seely, J. E., and Pegg, A. E. (1983) Ornithine decarboxylase (mouse kidney). Methods Enzymol 94, 158–161

63. Zhang, L., Mizumoto, K., Sato, N., Ogawa, T., Kusumoto, M., Niiyama, H., and Tanaka, M. (1999) Quantitative determination of apoptotic death in cultured human pancreatic cancer cells by propidium iodide and digitonin. Cancer Letters 142, 129–137

64. Vichai, V., and Kirtikara, K. (2006) Sulforhodamine B colorimetric assay for cytotoxicity screening. Nature Protocols 1, 1112–1116

